# Multivalent Ligand-Protein Interactions Using Polymeric Lysosome-Targeting Chimeras (PolyTACs) Leads to Lysosome-Targeting Receptor-Independent Degradation of Transmembrane Proteins

**DOI:** 10.1101/2025.05.06.652519

**Authors:** Ranit Dutta, Yasin Alp, Prachi Gupta, Badal Singh, S. Thayumanavan

## Abstract

Targeted protein degradation is growing rapidly as a therapeutic approach, with intracellular proteins degraded via the ubiquitin-proteasome or the autophagosome system and membrane proteins mainly through the lysosomal pathway. Current lysosomal degradation strategies rely on lysosome-targeting receptors (LTRs), limiting their applicability. We propose that multivalent non-covalent interactions on the cell membrane can drive lysosomal degradation of membrane proteins without the need of LTRs. To demonstrate this, we designed antibody-polymer conjugates, *viz*. Polymeric Lysosome-Targeting Chimeras (PolyTACs) functionalized with ligands that would non-covalently bind with transmembrane non-LTR proteins, *viz*., helper proteins on the cell surface in a polyvalent fashion. Cetuximab-based PolyTACs decorated with 4-(2-aminoethyl)benzenesulfonamide (ABS) ligands that cause multivalent interactions with membrane carbonic anhydrases induced degradation of EGFR, while atezolizumab- and trastuzumab-based PolyTACs effectively degraded PD-L1 and HER2, respectively. Additionally, PolyTACs using PD-L1 as the helper protein further improved degradation. Mechanistic studies confirmed clathrin- and caveolae-mediated endocytosis followed by lysosomal degradation of the target proteins. This LTR-independent nature of the approach offers opportunities for tissues targeting in membrane protein degradation that could open up new avenues in therapeutic strategies.

## Introduction

Targeted degradation of intra- and extracellular proteins has been growing rapidly as an area of interest both in academia and industry, owing to the fact that degradation of a protein-of-interest (POI) is a relatively permanent fate relative to occupancy-based inhibition.^1,2^ While intracellular POIs are mainly degraded by hijacking the ubiquitin-proteasome or the autophagasome machinery, membrane proteins and soluble extracellular proteins are often directed to the lysosome for proteolysis. To degrade these latter class of proteins, lysosome-targeting chimeras (LYTACs) that co-opt cation-independent mannose-6-phosphate as the lysosomal targeting receptor (LTR) were developed.^3,4^ LYTACs form a multicomponent complex with the LTR and POI, which causes the latter to be directed to the lysosome for degradation. Following this initial report, other versions of lysosomal protein degraders, such as bispecific aptamer chimeras, GlueTACs, and KineTACs have been developed.^5–7^ All these innovations involve the identification and leveraging of different types of LTRs. We envisaged that the reliance on LTRs can limit their utilization because many LTRs are found across various tissues, which could limit their ability to target specific diseased tissues effectively. Conversely, LTRs that are only present in certain tissues may not be suitable for targeting POIs in other tissue types. Therefore, by eliminating the need for LTRs, we could potentially increase the versatility of the lysosomal degradation system. We posit that simple multivalent interactions are sufficient to drive lysosomal degradation of POIs without relying on LTRs. In fact, we have shown that this can be achieved using a covalent multivalent interactions between disulfide functionalities on the cell surface and thiols on our custom-designed degraders.^8^ In this effort, we focus on greatly expanding the repertoire of this concept using non-covalent ligand-protein interactions. Reversibility and the tunability of these receptor-specific interactions on the cell surface offer to open up new opportunities. To test this idea here, we decorate polymer chains with multiple ligands targeting non-lysosome-targeting cell surface proteins and conjugate these polymers to a transmembrane POI-specific antibody. To distinguish these cell surface proteins from LTRs, we simply refer them as helper proteins. We term this non-covalent, ligand-decorated antibody-polymer conjugate platform Polymeric Lysosome-Targeting Chimeras (PolyTACs). The system is composed of two components: an antibody that targets the POI and ligand-decorated polymer chains for multivalent non-covalent interactions with a second cell surface helper protein (Scheme 1).

**Scheme 1.**
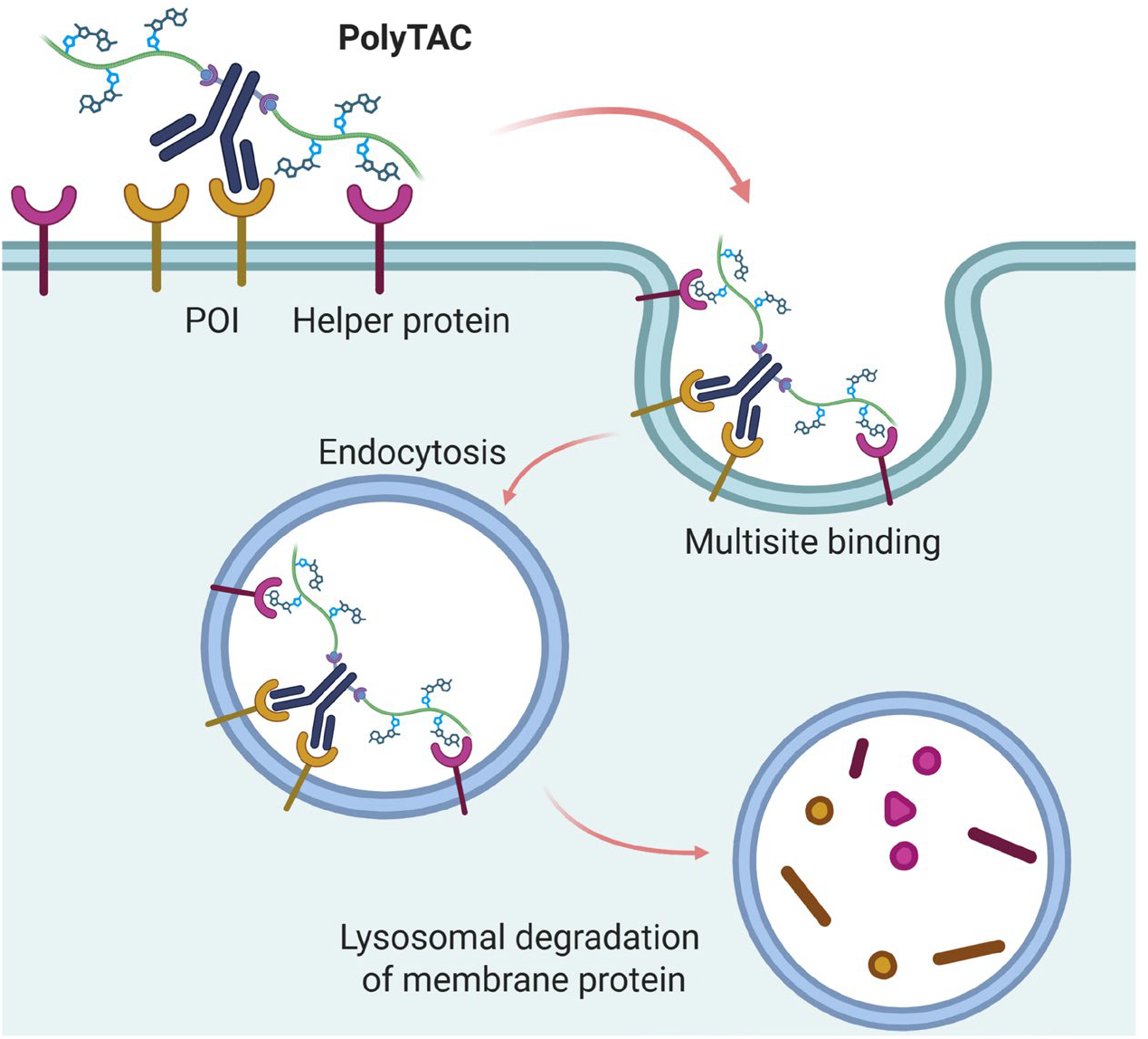
Schematic illustration of Polymeric Lysosome-Targeting Chimeras (PolyTACs) binding multivalently with transmembrane proteins and therefore driving the proteins toward lysosome for targeted membrane protein degradation.

## Results and Discussion

### 1. Synthesis of PolyTACs

To investigate the role of multisite interactions in driving lysosomal degradation, we designed out initial test case with an antibody-polymer conjugate featuring 4-(2-aminoethyl)benzenesulfonamide (ABS)^9^ moieties grafted onto the polymer side chains. We hypothesized that upon binding of the mAb to the POI, the ABS units on the polymer would engage with cell surface carbonic anhydrase isoforms, *viz*., CA-IX and CA-XII. This multivalent engagement is expected to promote receptor-mediated endocytosis and ultimately direct the POI to the lysosome.

To test this concept, we conjugated ABS-functionalized polymers to cetuximab (Ctx), an EGFR-targeting mAb, to generate Ctx-PolyTACs.^10^ EGFR is a transmembrane receptor implicated in cell proliferation and survival and is frequently overexpressed or mutated in various cancers.

For antibody-polymer conjugation, we employed an inverse electron demand Diels-Alder (IEDDA) reaction between trans-cyclooctene (TCO) and tetrazine (Figure 1a). First, Ctx was modified by incubation with TCO-PEG4-NHS ester (30 equivalents) for 2 h at room temperature, resulting in 10-15 TCO linkers per antibody, as assessed by LC-MS analysis (Figure 1b). In parallel, the ABS-functionalized polymer was prepared by first amidating tetrazine-PEG4-amine with the carboxylic acid terminus of a PEG-r-co-azide polymer (M_n_ ∼15 kDa, 40 PEG6 units, 10 azides) followed by installation of the ABS ligands onto the polymer side chains. DBCO-PEG8-amine functionalized with ABS was subjected to a strain-promoted azide-alkyne cycloaddition (SPAAC) reaction with azide-functionalized polymer, yielding ∼8 ABS units per polymer chain. Importantly, this approach preserved orthogonality with the tetrazine handle on the chain end, allowing modular ligand incorporation without altering the polymer backbone.

**Figure 1:**
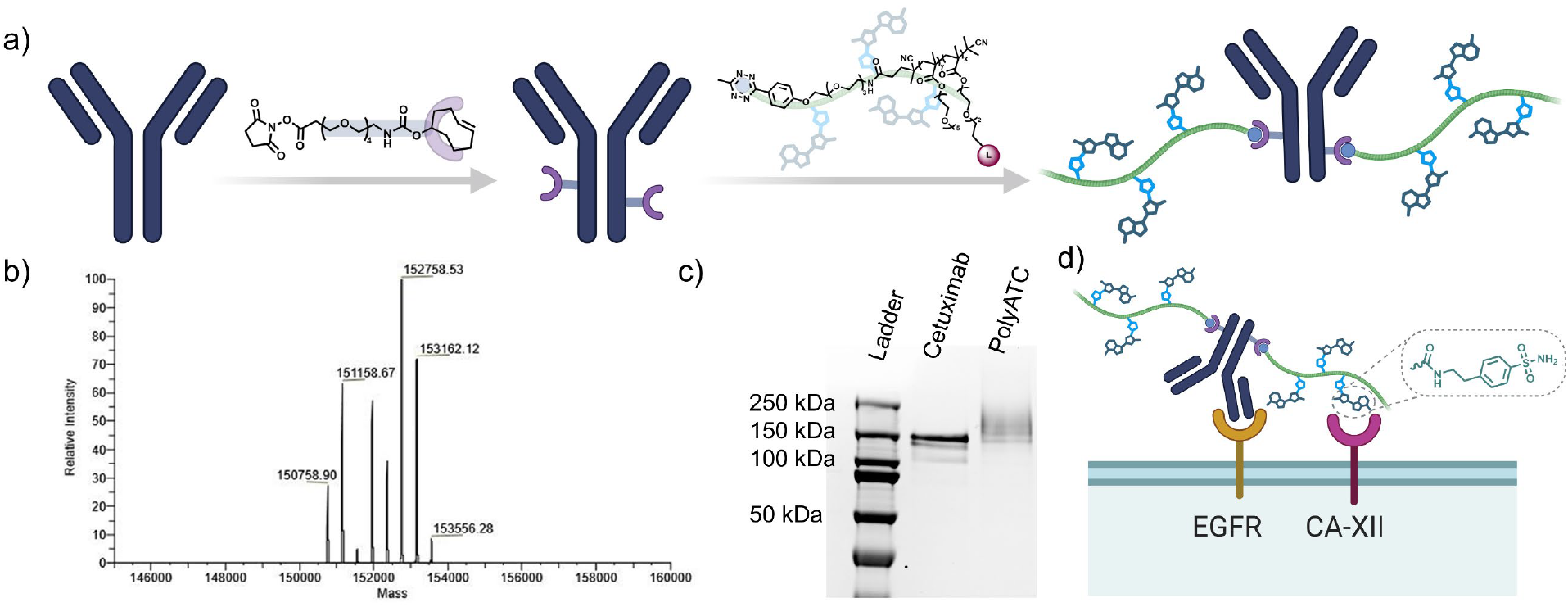
(a) Schematic illustration of synthesis of PolyTAC in two steps: TCO-PEGn-NHS ester conjugation with monoclonal antibody (mAb) followed by TCO-tetrazine conjugation with ligand-decorated polymer. (b) LC-MS results showing degree of TCO conjugation with mAb. (c) SDS-PAGE result showing complete consumption of mAb after TCO-tetrazine bioconjugation. (d) Schematic illustration of Ctx-4-PolyTAC simultaneously binding with EGFR and CA-XII on cell surface.

Finally, the TCO-modified Ctx was reacted with 30 equivalents of the tetrazine-functionalized ABS polymer (ABS-pol) at room temperature for 2 h, resulting in quantitative antibody consumption as assessed by SDS-PAGE (Figure 1c), to yield the final construct: Ctx-ABS-PolyTAC (Ctx-4-PolyTAC) (Figure 1d). Here “4” denotes the PEG4 linker connecting the antibody to the TCO and consequently to the polymer scaffold.

### 2. Structural optimization and degradation efficacy of PolyTACs

We began evaluating the efficacy of PolyTACs for membrane protein degradation by treating the EGFR-overexpressing triple-negative breast cancer cell line MDA-MB-231 with Ctx-4-PolyTAC for 24 h.

Compared to PBS and cetuximab (Ctx) controls, we observed a significant dose-dependent reduction in EGFR levels with ABS-pol, with both the half-maximal degradation concentration (DC_50_) and maximum degradation percentage (D_max_, ∼50%) achieved at ∼1 µM, as assessed by western blot (Figure 2a,b). To complement these results and more quantitatively assess surface EGFR levels, we also employed flow cytometry. Cells were treated with PolyTAC for 24 h, followed by PBS washes and incubation with Alexa Fluor 488-labeled anti-EGFR antibody (matuzumab that is known to bind a different epitope on EGFR than Ctx)^11^ for 1 h at 4 °C. After additional washing, surface EGFR expression was analyzed by flow cytometry. Consistent with the western blot data, treatment with 1 µM Ctx-4-PolyTAC led to ∼50% reduction in surface EGFR levels compared to controls (Figure 2c). Together, these findings demonstrate that multivalent PolyTACs can drive degradation of a POI without requiring an LTR. However, the degradation efficiency was modest compared to the reported LTR-targeted strategies, indicating the need for further optimization. We identified three potential avenues for improving degradation and for testing the broad applicability of the concept: (i) increasing the linker length between the antibody and polymer to enhance the flexibility and the ligand-grafted polymer’s reach toward the helper proteins after POI engagement; (ii) altering the helper protein targeted by the polymer-conjugated ligands (*e*.*g*., ABS for CA-IX/XII vs. BMS-8 for PD-L1); and (iii) improving ligand’s binding affinity by carefully varying the choice-of-ligand (*e*.*g*., BMS-8 vs. BMS-1166, both targeting PD-L1).

**Figure 2:**
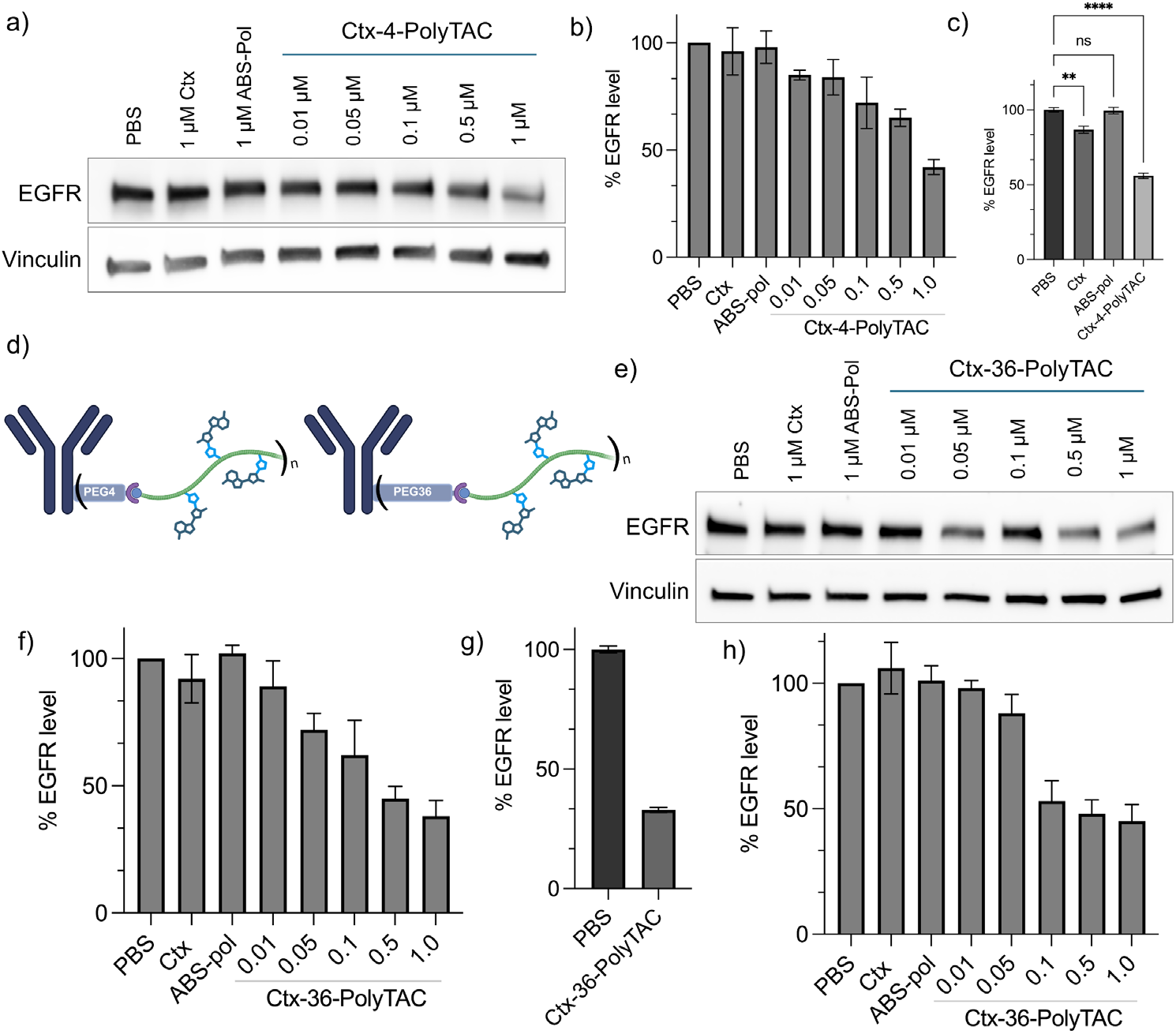
(a,b) Western blot results showing degradation percentage of EGFR after treating Ctx-4-PolyTAC in MDA-MB-231 cells for 24 h. (c) Flow cytometric analysis of percentage EGFR degradation after treating 1 µM solution of Ctx-4-PolyTAC in MDA-MB-231 cells for 24 h and comparison with the control treatments: PBS, 1 µM Ctx and 1 µM ABS-pol. (d) Schematic illustration showing variation of TCO-mAb linker length: 4 vs 36 PEG repeat units. (e,f) Western blot results showing degradation percentage of EGFR after treating Ctx-36-PolyTAC for 24 h in MDA-MB-231 cells. (g) Flow cytometric analysis of percentage EGFR degradation after treating 1 µM solution of Ctx-36-PolyTAC in MDA-MB-231 cells for 24 h and comparison with the control treatment: PBS. (h) Western blot results showing degradation percentage of EGFR by treating Ctx-36-PolyTAC in SK-BR-3 cells for 24 h.

First, to extend the length of the linker, we substituted the PEG4 linker with a longer PEG36 linker to generate Ctx-36-PolyTAC (Figure 2d). We hypothesized that upon EGFR binding, the POI-PolyTAC complex would occupy a substantial area on the cell surface, potentially limiting access to nearby helper membrane proteins like CA-IX/XII. By extending the linker length, we expect to improve the likelihood of multivalent interactions between the polymer and helper proteins. When MDA-MB-231 cells were treated with Ctx-36-PolyTAC under similar conditions, we observed a shift in DC_50_ to approximately 500 nM, while D_max_ was improved to 65% at 1 µM (Figure 2e,f,g). These results indicate that both polymer length and flexibility significantly influence degrader performance.

Additionally, the expression level of the helper protein is a critical factor for efficient multivalent engagement. For instance, CA-XII expression is lower in MDA-MB-231 cells compared to SK-BR-3 cells, which also overexpress EGFR. To explore this, we treated SK-BR-3 cells with Ctx-36-PolyTAC across the same concentration range. This resulted in improved EGFR degradation profile (DC_50_ between 100 and 500 nM, D_max_ ∼60% at 1 µM), confirming that PolyTAC efficacy depends on the expression levels of both POI and helper protein (Figure 2g). These findings support an “AND”-gated degradation mechanism, in which enhanced activity is achieved in cells that overexpress both target protein and the helper proteins. This dual-dependency strategy holds promise for achieving tissue-specific therapeutic effects while minimizing off-target degradation.

### 3. Investigating the modularity of the PolyTAC strategy

To validate our findings and evaluate the broader applicability of the strategy, we extended the chain length optimization approach to another therapeutically relevant membrane protein, Programmed Death-Ligand 1 (PD-L1). PD-L1 is an immune checkpoint protein that binds to PD-1 receptors on activated T and B cells, suppressing immune response against cancer cells. To target PD-L1, we used the FDA-approved monoclonal antibody atezolizumab (Atz),^12^ modified with both PEG4 and PEG36 linkers. Subsequent conjugation with the ABS-functionalized polymer yielded two PolyTAC variants: Atz-4-PolyTAC and Atz-36-PolyTAC.

We treated MDA-MB-231 cells with these PolyTACs for 24 h and assessed PD-L1 levels using both western blot and flow cytometry. Western blot analysis revealed an improved DC_50_ for Atz-36-PolyTAC compared to Atz-4-PolyTAC, indicating enhanced degradation efficiency with increased linker length (Figure 3a,b,c,d). Flow cytometry further confirmed a greater reduction in PD-L1 surface expression with Atz-36-PolyTAC (Figure 3e). To determine if degradation efficacy could be further improved with extended treatment time, we conducted a time-course analysis (6 h, 24 h, and 48 h). For both Ctx-36-PolyTAC and Atz-36-PolyTAC, we observed a time-dependent increase in POIs degradation (Figure 3f).

**Figure 3:**
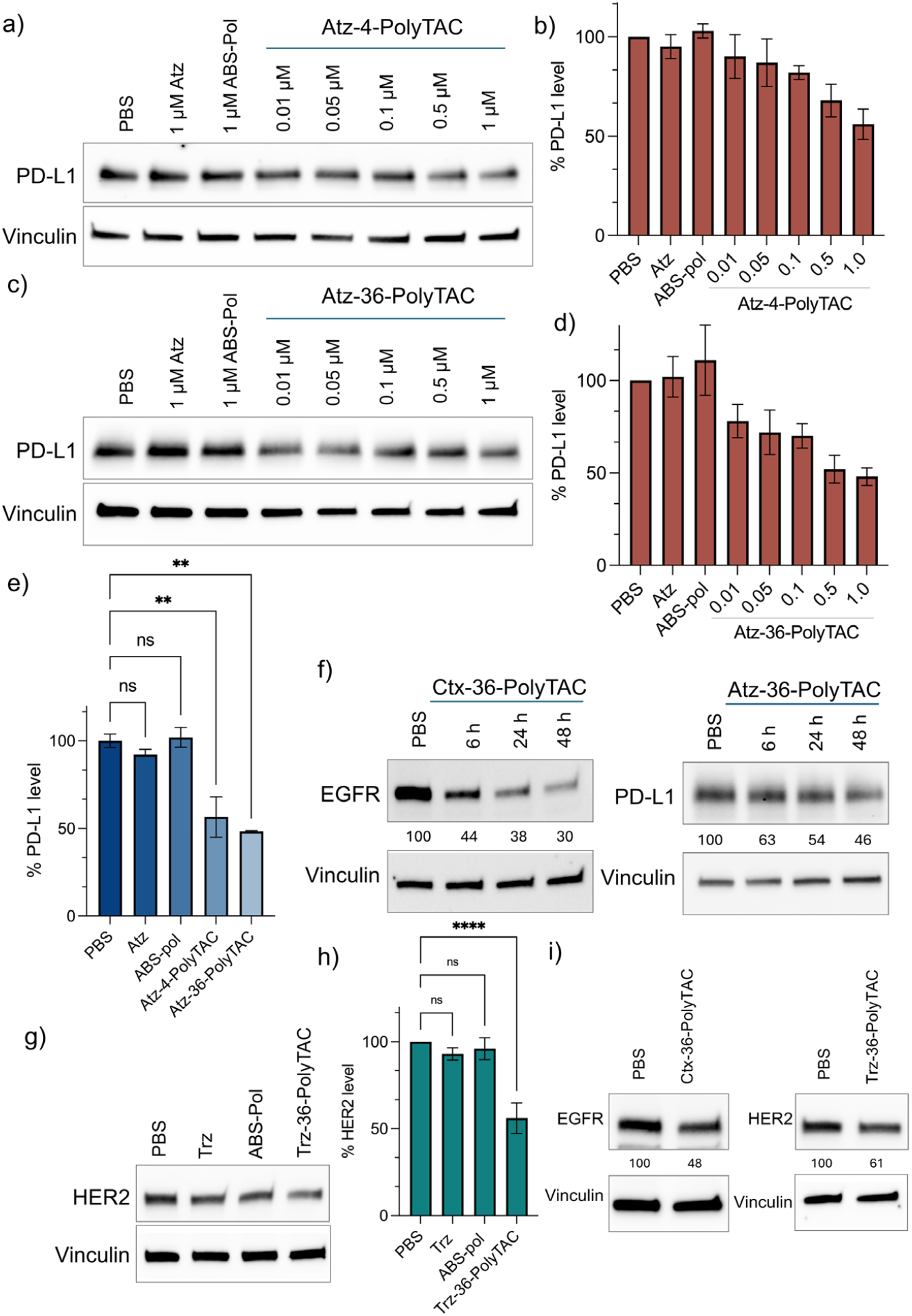
(a,b) Western blot results showing degradation percentage of PD-L1 after treating Atz-4-PolyTAC in MDA-MB-231 cells for 24 h. (c,d) Western blot results showing degradation percentage of PD-L1 after treating Atz-36-PolyTAC for 24 h in MDA-MB-231 cells. (e) Flow cytometric analysis of percentage PD-L1 degradation after treating 1 µM solutions of Atz-4-PolyTAC and Atz-36-PolyTAC in MDA-MB-231 cells for 24 h and comparison with the control treatments: PBS, 1 µM Atz and 1 µM ABS-pol. (f) Western blot analysis of EGFR and PD-L1 degradation using Ctx-36-PolyTAC and Atz-36-PolyTAC respectively for 6, 24 and 48 h treatments. (g,h) Western blot results showing degradation percentage of HER2 by treating Trz-36-PolyTAC in SK-BR-3 cells for 24 h. (i) Western blot results showing degradation of EGFR and HER2 after cotreating 1 µM solutions of Ctx-36-PolyTAC and Trz-36-PolyTAC in SK-BR-3 cells for 24 h.

We next targeted another clinically significant membrane protein, human epidermal growth factor receptor 2 (HER2), which is overexpressed in aggressive breast cancers and associated with rapid tumor progression. By conjugating trastuzumab (Trz)^13^ with ABS-polymer via PEG36 linkers, we generated Trz-36-PolyTAC and treated HER2+ SK-BR-3 cells for 24 h. This led to ∼45% degradation of HER2, further supporting the broad applicability of our modular platform (Figure 3g,h). This strategy also allows for combinatorial elimination of multiple POIs. Given that co-targeting EGFR and HER2 using cetuximab and trastuzumab has shown synergistic effects in reducing tumor hypoxia *in vivo*^14^, we explored a dual-degradation strategy by treating SK-BR-3 cells with a combination of Ctx-36-PolyTAC and Trz-36-PolyTAC for 24 h. This resulted in potent degradation of both EGFR and HER2, comparable to single PolyTAC treatments (Figure 3i). These findings highlight the potential of PolyTACs for combination therapy, particularly in contexts where monotherapy is insufficient or prone to resistance emergence.

### 4. Helper protein selection and ligand affinity govern the efficiency and scope of PolyTAC-mediated membrane protein degradation

One limitation of using carbonic anhydrase isoform CA-XII as helper protein is its ubiquitous expression across healthy and cancerous cell lines. Moreover, when we immunoblotted CA-XII for the treatments of Ctx-36-PolyTAC and Atz-36-PolyTAC, we observed insignificant degradation of CA-XII at different concentrations (Figure 4a,b). Ctx-36-PolyTAC treatment in the CA-XII overexpressing SK-BR-3 cell line also showed only nominal degradation of CA-XII (Figure 4c). This limited susceptibility may stem from its high translation rate and rapid endosomal recycling, likely due to its essential physiological roles in pH regulation and fluid balance.^15^ Consistent with this observation, CellTiter-Glo assays performed after 48 h PolyTAC treatments showed cell viabilities comparable to those observed with Ctx and Atz treatments alone, suggesting that insufficient CA depletion contributes to the lack of cytotoxic effects (Figure 4d).

**Figure 4:**
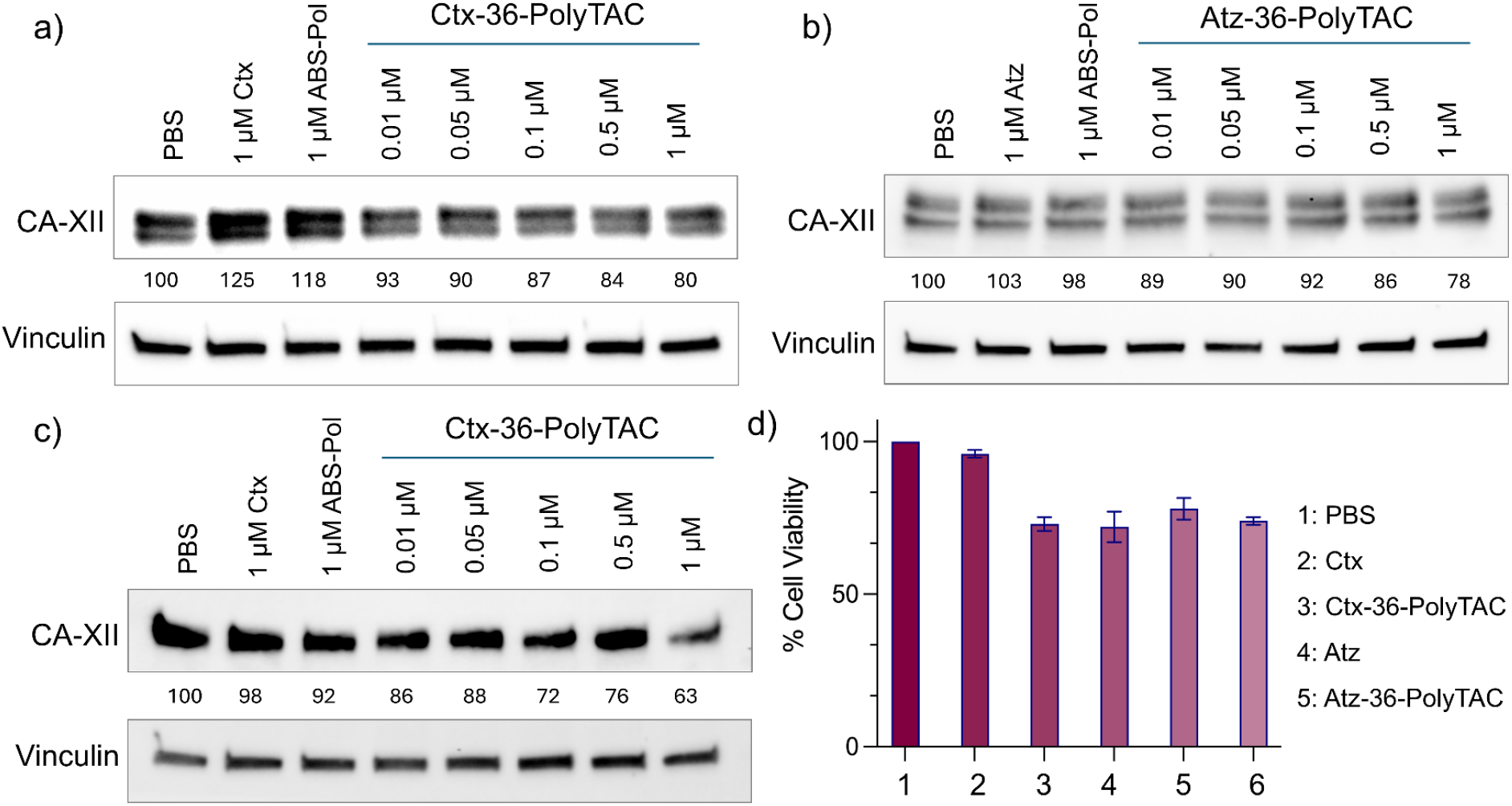
(a) Western blot results showing degradation percentage of CA-XII by treating Ctx-36-PolyTAC in MDA-MB-231 cells for 24 h. (b) Western blot results showing degradation percentage of CA-XII by treating Atz-36-PolyTAC in MDA-MB-231 cells for 24 h. (c) Western blot results showing degradation percentage of CA-XII by treating Ctx-36-PolyTAC in SK-BR-3 cells for 24 h. (d) CellTitre-Glo assay showing percentage cell viability after treating Ctx (1 µM), Ctx-36-PolyTAC (1 µM), Atz (1 µM) and Atz-36-PolyTAC (1 µM) for 48 h in MDA-MB-231 cells.

To test if degradation of POI can be achieved using the ligand-protein interaction in the polymer in the antibody-polymer conjugate, we pivoted to targeting PD-L1 as the helper protein, as our Atz-based studies shows that PD-L1 can be degraded using the antibody targeting. We functionalized Ctx with polymer chains bearing BMS-8, a small molecule ligand targeting the extracellular domain of PD-L1^16^. Given that MDA-MB-231 cells co-express EGFR and PD-L1, treatment with the resulting Ctx-BMS8-PolyTAC for 24 h led to the degradation of both EGFR and PD-L1 (Figure 5a,b).

**Figure 5:**
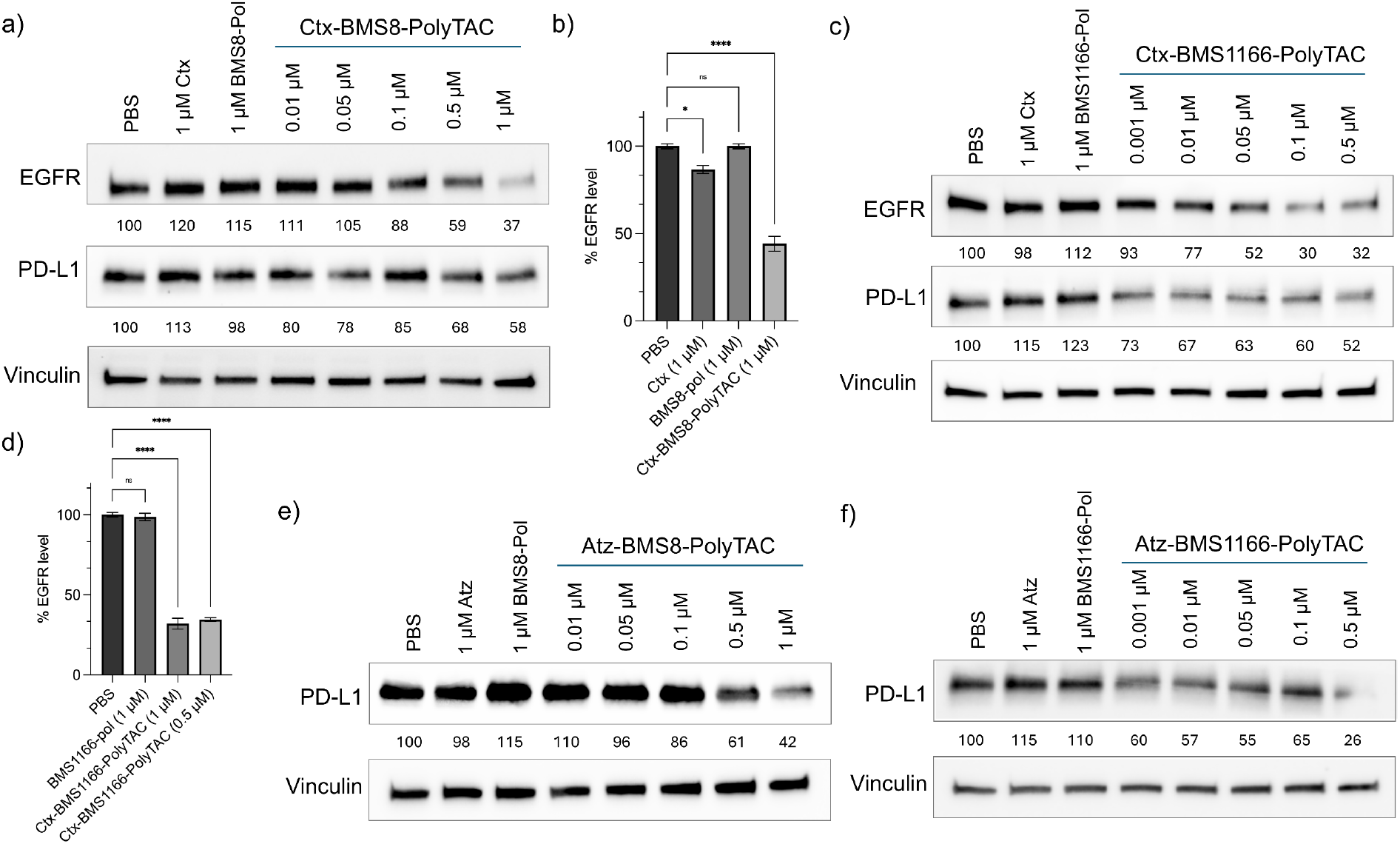
(a) Western blot results showing degradation percentage of EGFR and PD-L1 after treating Ctx-BMS8-PolyTAC in MDA-MB-231 cells for 24 h. (b) Flow cytometric analysis of percentage EGFR degradation after treating 1 µM solutions of Ctx-BMS8-PolyTAC in MDA-MB-231 cells for 24 h and comparison with the control treatments: PBS, 1 µM Ctx and 1 µM BMS8-pol. (c) Western blot results showing degradation percentage of EGFR and PD-L1 after treating Ctx-BMS1166-PolyTAC in MDA-MB-231 cells for 24 h. (d) Flow cytometric analysis of percentage EGFR degradation after treating 0.5 µM solutions of Ctx-BMS1166-PolyTAC in MDA-MB-231 cells for 24 h and comparison with the control treatments: PBS, 0.5 µM Ctx and 0.5 µM BMS1166-pol. (e) Western blot results showing degradation percentage of PD-L1 after treating Atz-BMS8-PolyTAC in MDA-MB-231 cells for 24 h. (f) Western blot results showing degradation percentage of PD-L1 after treating Atz-BMS1166-PolyTAC in MDA-MB-231 cells for 24 h.

We further hypothesized that using higher affinity ligands could enhance degradation efficiency. To this end, we replaced BMS-8 with BMS-1166, a PD-L1 ligand with ∼100-fold better binding affinity.^16^ BMS-1166 polymers were conjugated with Ctx to generate Ctx-BMS1166-PolyTAC. As anticipated, these PolyTACs exhibited substantially improved potency. Ctx-BMS1166-PolyTAC showed DC_50_ value of ∼50 nM, with D_max_ values reaching ∼70% at 500 nM (Figure 5c,d), compared to Ctx-BMS8-PolyTAC’s DC_50_ and D_max_ of ∼700 nM and ∼60% respectively. Flow cytometry further corroborated these findings, showing effective degradation of EGFR. Furthermore, Ctx-BMS1166-PolyTAC induced enhanced degradation of the helper protein PD-L1 compared to the Ctx-BMS8 conjugate. This highlights a key design principle: by combining high-affinity ligands with carefully selected helper proteins, PolyTACs can be engineered to achieve efficient, multiprotein degradation with a single molecular construct.

Next, we explored the concept of homofunctional PolyTACs, designed to target a single protein using both the antibody and the ligands by conjugating the BMS-8 and BMS-1166 ligands containing polymers to Atz, resulting in Atz-BMS8-PolyTAC and Atz-BMS1166-PolyTAC respectively. These homo-PolyTACs led to significant degradation of PD-L1 (Figure 5e,f). As seen for the Ctx conjugates, the Atz-BMS1166-PolyTAC showed higher efficacy compared to Atz-BMS8-PolyTAC.

### 5. Mechanistic insights into endocytic and lysosomal processing of PolyTAC-targeted proteins

To elucidate the internalization and degradation mechanisms underlying PolyTAC-induced protein degradation, we employed different strategies to decipher (*a*) the role of multivalent interactions offered by the ligand-decorated polymer chains in causing protein degradation, and (*b*) the underlying endocytic/lysosomal pathways by utilizing a panel of inhibitors. Firstly, we co-treated MDA-MB-231 cells with the ABS-pol (100 µM) or ethoxzolamide^9^ (EZA, 100 µM), a comparatively stronger inhibitor of CAs, along with Ctx-36-PolyTAC for 24 h. We also treated cells with Ctx conjugated with polymer that lacks the ABS ligands (Ctx-pol, 1 µM) separately. In all cases, we observed no degradation of EGFR, validating the role of multivalency in degradation (Figure 6a).

**Figure 6:**
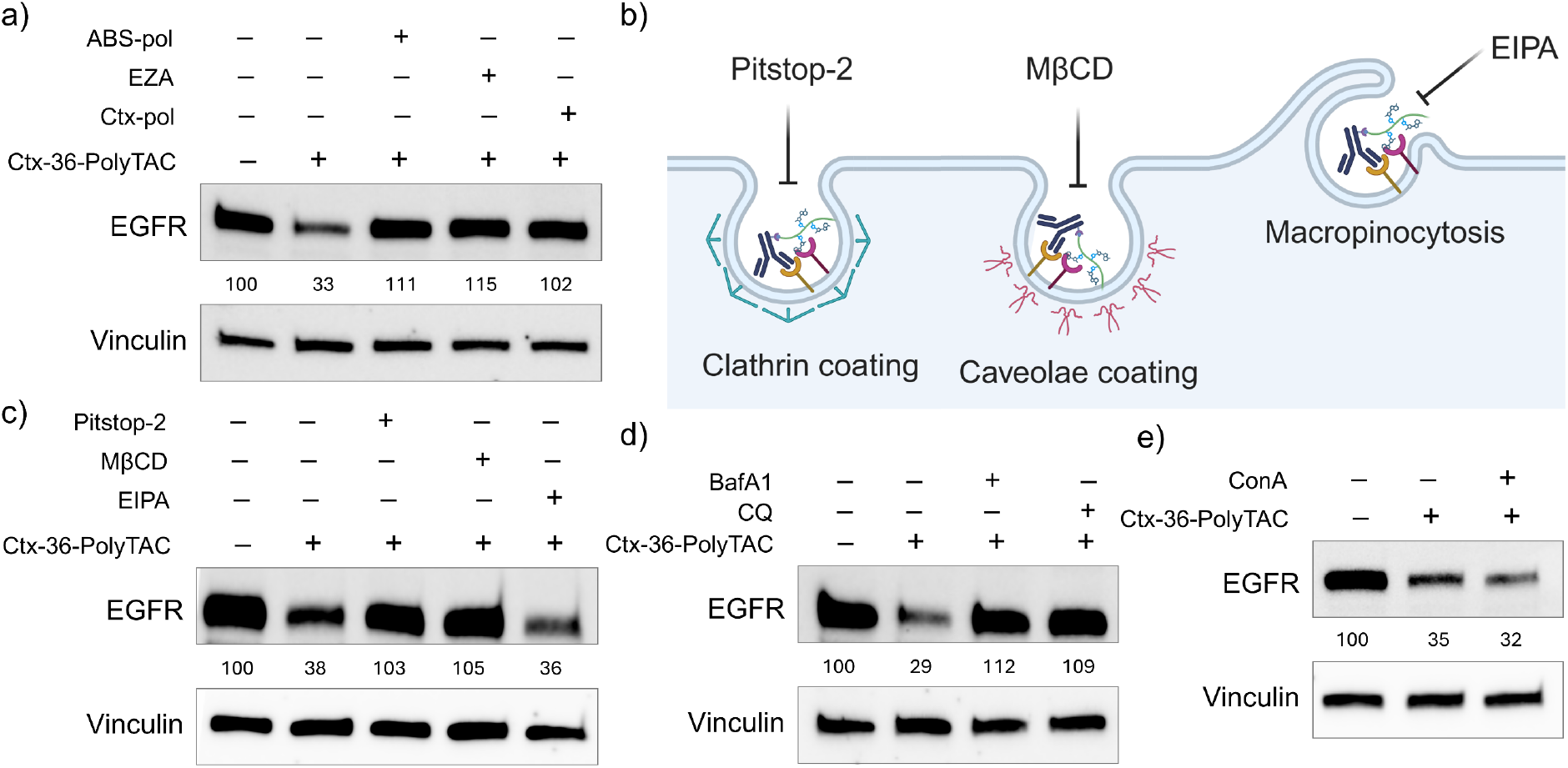
(a) Investigating the role of multivalent binding of the PolyTAC with POI and the helper protein in determining the degradation efficacy by utilizing specific inhibitors: ABS-pol and ethoxzolamide (EZA) blocking CA-XII and Ctx-pol blocking EGFR when cotreated with Ctx-36-PolyTAC. (b) Schematic illustration of endocytosis inhibitors blocking clathrin and caveolae-dependent pathways and macropinocytosis. (c) Western blot analysis of EGFR degradation: MDA-MB-231 cells were pretreated with clathrin- and caveolae-dependent endocytosis and macropinocytosis inhibitors (20 µM pitstop-2, 0.5 mM MβCD and 10 µM EIPA respectively) for 30 min followed by washing and treatment of Ctx-36-PolyTAC for 24 h. (d) Western blot analysis of EGFR degradation: MDA-MB-231 cells were cotreated with endolysosomal degradation inhibitors (100 nM BafA1 and 100 µM CQ) and Ctx-36-PolyTAC for 24 h. (e) Western blot analysis of EGFR degradation: MDA-MB-231 cells were cotreated with autolysosomal degradation inhibitor (10 nM ConA) and Ctx-36-PolyTAC for 24 h.

Then we sought to identify the modes of cellular uptake for the POI-PolyTAC-helper protein complex. MDA-MB-231 cells were pretreated for 30 minutes with inhibitors of specific endocytic pathways: Pitstop-2 (a clathrin-mediated endocytosis inhibitor, 20 µM), methyl-β-cyclodextrin (MβCD, a caveolae-mediated endocytosis inhibitor, 0.5 mM), and 5-(*N*-ethyl-*N*-isopropyl)-amiloride (EIPA, a macropinocytosis inhibitor, 10 µM)^17–19^ (Figure 6b). After pretreatment, the cells were washed and subsequently treated with either Ctx-36-PolyTAC or Atz-36-PolyTAC for 24 h. Western blot analysis revealed that both Pitstop-2 and MβCD pretreatments resulted in a near-complete inhibition of POI degradation, whereas EIPA had no significant effect (Figure 6c). These results suggest that internalization of the PolyTAC constructs occurs predominantly through clathrin- and caveolae-mediated endocytosis, while macropinocytosis does not play a role in the uptake of the complex.

Next, we investigated the intracellular degradation machinery responsible for processing the internalized complexes. To determine whether the lysosomal pathway is indeed involved, cells were pretreated with bafilomycin A1 (BafA1, 100 nM), an inhibitor of vacuolar H^+^-ATPase that blocks lysosomal acidification, and chloroquine (CQ, 100 µM), a lysosomotropic agent that impairs endosome acidification.^20^ Both BafA1 and CQ pretreatments led to a substantial suppression of POI degradation, confirming that the PolyTAC-mediated process is reliant on lysosomal processing (Figure 6d). We also treated cells with concanamycin A (ConA, 10 nM), an inhibitor of autolysosomal acidification^20^, and still observed robust POI degradation (Figure 6e). This result rules out the role of autolysosomes in the PolyTAC-mediated degradation pathway. Together, these findings establish that PolyTAC internalization is primarily mediated via clathrin- and caveolae-dependent endocytosis, and that the subsequent degradation of the target protein occurs through the endolysosomal pathway.

## Conclusion

In summary, we present a modular strategy for transmembrane protein degradation using PolyTAC platform, which harnesses multivalent non-covalent interactions to drive lysosomal processing of membrane-bound targets. By integrating multi-ligand decorated polymers with monoclonal antibodies, we successfully engineered a series of PolyTACs capable of selectively degrading POIs without relying on LTRs. Our study highlights several key principles for optimizing the design and functionality of PolyTACs: (*i*) Versatility across targets: PolyTACs effectively degraded various POIs when paired with different combinations of monoclonal antibodies and non-covalent ligands, illustrating the broad applicability of this platform. (*ii*) LTR-independent internalization: We demonstrated that non-LTR transmembrane proteins can serve as helper proteins to drive multivalency-induced internalization and subsequent degradation of POIs, expanding the range of potential targets. (*iii*) Structural considerations for efficacy: The design of potent PolyTACs requires the incorporation of long, flexible linkers between the antibody and the polymer backbone, as well as ligands with high binding affinity. (*iv*) Contextual specificity: Effective degradation was observed in cell lines where both the POI and helper protein were co-expressed, enabling context-dependent and potentially tissue-selective degradation. (*v*) Dual-targeting potential: Depending on the nature of the helper protein and the ligands used, PolyTACs can be tuned to degrade both the POI and the helper protein simultaneously, opening up new avenues for multi-protein knockdown using a single conjugate.

While the platform shows significant promise, further optimization is required, particularly in refining polymer architecture, reducing the overall molecular weight of the construct, enhancing biocompatibility, and achieving a favorable pharmacokinetic profile. In vivo validation and translational development of these degraders form a major focus of our ongoing efforts. Taken together, our PolyTAC approach introduces a novel, LTR-independent strategy for targeted extracellular protein degradation, offering a versatile and potentially tissue-specific therapeutic modality.

